# A pili-driven bacterial turbine

**DOI:** 10.1101/2022.02.14.480354

**Authors:** Wolfram Pönisch, Vasily Zaburdaev

## Abstract

Work generated by self-propelled bacteria can be harnessed with the help of microdevices. Such nanofabricated microdevices, immersed in a bacterial bath, may exhibit unidirectional rotational or translational motion. Swimming bacteria that propel with the help of actively rotating flagella are a prototypical example of active agents that can power such microdevices.

In this work, we propose a computational model of a micron-sized turbine powered by bacteria that rely on active type IV pili appendages for surface-associated motility. We find that the turbine can rotate persistently over a time scale that significantly exceeds the characteristic times of the single cell motility. The persistent rotation is explained by the collective dynamics of multiple pili of groups of cells attaching to and pulling on turbine. Furthermore, we show that the turbine can rotate permanently in the same direction by altering the pili binding to the turbine surface in an asymmetric fashion. We thus can show that by changing the adhesive properties of the turbine while keeping its symmetric geometry, we can still break the symmetry of its rotation.

Altogether, this study widely expands the range of bacteria that can be used to power nanofabricated microdevices, and, due to high pili forces generated by pili retraction, promises to push the harnessed work by several orders of magnitude.

## 1 INTRODUCTION

The last years have seen significant advances in nanofabrication, permitting the invention of a wide range of micron-sized artificial devices. A fascinating question is how such devices can be powered by actively moving biological matter, typically consisting of bacteria. Examples are beads that move due to collision or attachment of cells to its surface [1, 2], swimming devices due to bacterial carpets attached to their surface [3], and geometrically asymmetric devices immersed in a bath of actively moving cells that rotate due to random collisions with the cells [4, 5, 6, 7]. Usually, bacteria that swim with the help of rotating flagella were employed. In this study, we propose to consider twitching bacteria that exhibit surface-associated locomotion mediated by type IV pili [8, 9]. Pili are microns long active polymeric appendages protruding from the cell membrane. They undergo cycles of protrusion and retraction and can bind to a substrate and the pili of other cells. The combination of these two processes leads to aggregation of cells [10, 11, 12] and twitching motility on a substrate [13, 14, 15, 16] by a mechanism reminiscent of a grappling hook (see Fig. 1A).

**Figure 1.**
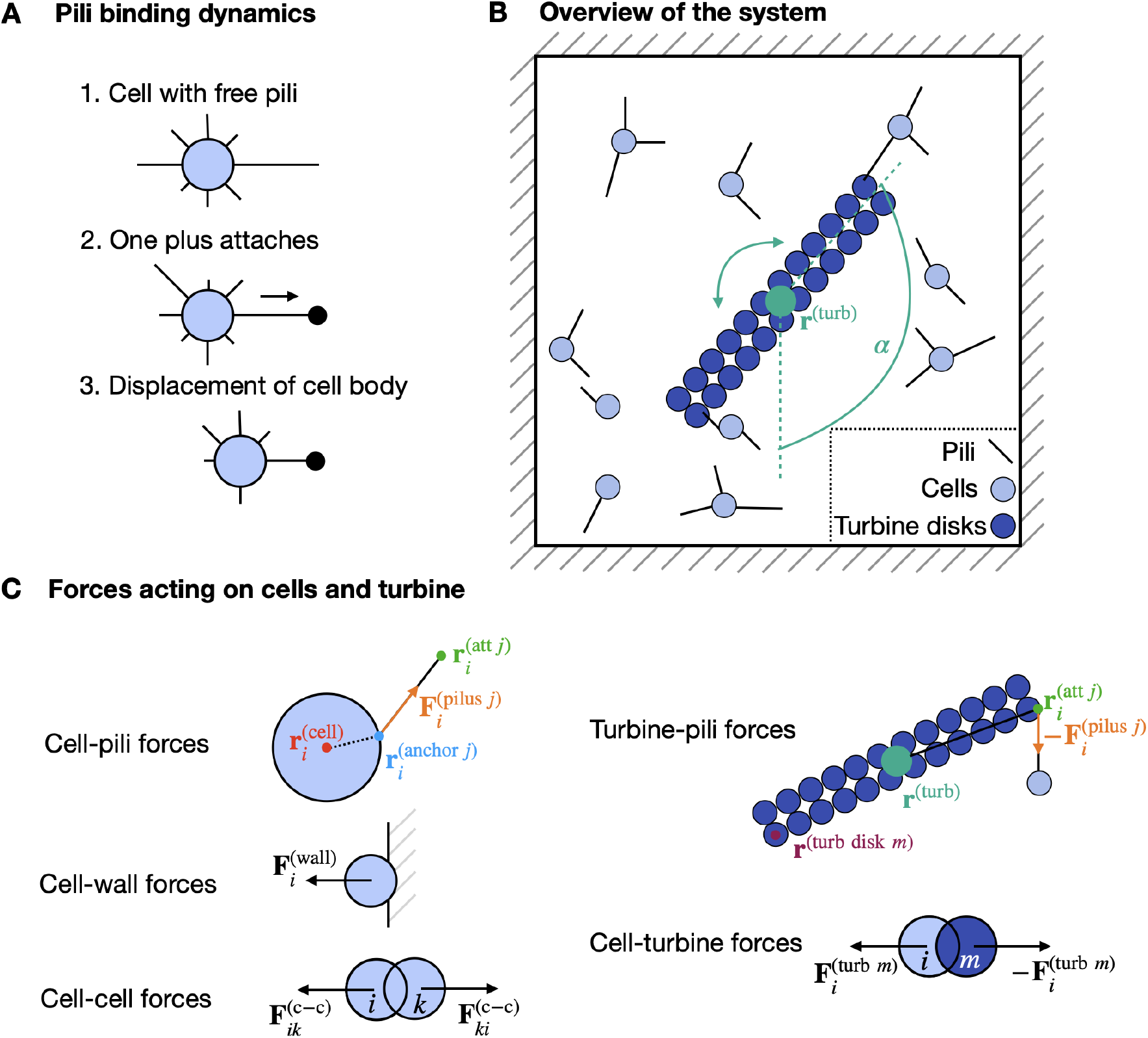
Overview of the computational model of bacteria driving a rotatable turbine by attachment of type IV pili. **(A)** Sketch of how pili binding and retraction can lead to a force acting on a cell, a process reminiscent of a grappling hook. **(B)** Overview of the system. Cells are located in a box together with the rotatable turbine. The cells possess pili which they use for the motility on the substrate and for attachment to the turbine. The turbine center is located at **r**^(turb)^ and has an orientation defined by the angle *α*. **(C)** Summary of all forces acting on the cells and turbine. Friction forces with the environment (surrounding liquid and substrate) are not shown.

Here, we investigate how bacteria that can attach type IV pili to a micron-sized turbine can drive its rotation. This system is especially interesting since multiple pili belonging to an individual bacterium can generate total forces in the range of nano-Newtons [17]. In contrast to that, bacteria that swim with the help of flagella typically have thrust forces in the order of 1-10 pN [18]. The molecular motor involved in the disassembly an hence, in the retraction of an individual pilus, called pilT, can generate forces in the range of 100 − 180 pN [19]. This makes pilT the strongest molecular motor known in nature, with forces 2-20 times larger than those generated by kinesins or polymerases [17]. Hence, we are asking if cells with type IV pili that can create so large forces might be more attractive candidates to power microdevices.

In this work, by means of a computational model, we study the dynamics of a rotatable turbine immersed in a bath of twitching bacteria. We investigate how the turbine rotation is affected by the binding and unbinding of the bacterial pili. We find that, due to the adhesion of multiple pili to the turbine over time scales that can strongly exceed the characteristic time scales of the individual pili attachment, the turbine can persistently rotate in one direction over extensive durations. While these persistent rotations have a limited lifetime, one can engineer a system where they become permanently unidirectional by introducing asymmetric binding of the pili to the turbine.

## 2 MATERIALS AND METHODS

First, we introduce the computational model of twitching bacteria and their interactions with a turbine. A related version of this computational model was considered previously to describe bacterial surface motility [16] and bacterial aggregates [20, 21, 22].

While we consider the bacterium *Neisseria gonorrhoeae* as the primary biological example, the computational model can be easily adapted to account for other bacteria that use type IV pili, e.g. *Neisseria meningitidis* [11] or *Pseudomonas aeruginosa* [23]. In the following, we focus on the regime of low cell density. This allows us not to consider cell-cell interactions and the formation of bacterial aggregates mediated by the binding of the pili of different cells. As a result, we do not expect three dimensional aggregates to form [20, 21]. This also enables us to only consider a simpler two-dimensional system.

### 2.1 Geometry of the cells, pili and turbine

The experimental system we are mimicking are bacterial cells confined in a box with the turbine at its center (see Fig. 1B). The cells can move over the substrate via pili binding and unbinding and additionally interact with the turbine by pili attachment and excluded volume effects.

While *Neisseria gonorrhoeae* cells typically have a diplococcus shape [24], we simplify the *in silico* cell shape of bacteria as two-dimensional circular disks with a radius *R*. Please note, however, that a diplococcus shape can be also considered [16, 20, 21, 22]. Each cell possesses exactly *N*_p_ pili that are homogeneously distributed on the cell outline (see Fig. 1A). We approximate pili as straight lines connecting two points: their start point (also called anchor point), located at the circular cell surface, and their end point. The distance between these two points is called the contour length *l*_c_ of the pilus.

The turbine is described by *N*_turb_ × 2 disks (with radius *R*) arranged in a double row array (see Fig. 1B). Since the only way how the turbine can move is by rotation, the relative locations of the turbine disks towards each other are fixed. The orientation of the turbine is described by the angle *α* with respect to the *y*-axis and the turbine center is located at the position **r**^(turb)^ (see Fig. 1B).

### 2.2 Pili dynamics and binding properties

Initially, a pilus protrudes with the velocity *v*_p_ in a direction perpendicular to the cell surface. When a pilus reaches a specific length, drawn from an exponential length distribution with mean length *l*_p_ [15, 14], it starts to retract with a velocity 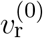. This retraction continues until the pilus has a contour length *l*_c_ = 0. In that case, the pilus is removed and immediately, a new pilus protrudes from the cell from the same position.

A pilus binds stochastically to the substrate or the turbine disks, independently whether it is protruding or retracting. In both cases, a pilus can only bind with its tip and it can bind only to either the substrate or the turbine disks. After binding, a pilus immediately starts to retract [25]. The binding is modeled as a Poisson process with the binding rate 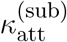 to the substrate and 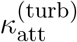 to any of the turbine disks. Since the model is two-dimensional, we ignore that, for pili to bind to a substrate, they first need to be long enough to reach the substrate with their tips. In 3D, this leads to a delay in the initial binding event of a newly protruding pilus, but is less relevant for following attachments of pili. Hence, initially, the binding rate might be smaller to allow a pilus to protrude to reach the substrate. To account for this process, pili that bind to the substrate the first time do so with a rate 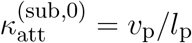. Again, the binding is modelled as a Poisson process. The rate 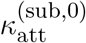 corresponds to the binding of a pilus that protrude with the velocity *v*_p_ until it reaches a length determined by an exponential distribution with mean length *l*_p_, in which case it binds. On average, this takes the time 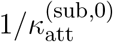.

An attached pilus will generate a pulling force that acts on the cell and, if attached to a turbine disk, also on the turbine. Each pilus is modeled as a Hookean spring with the spring constant *k*_p_. After attachment, a pilus is stretched due to its retraction and hence, mediates a pulling force. This force is proportional to the difference between the contour length *l*_c_ and the length of the pilus that it would have if it was not attached, here called the free length *l*_f_. Here, we consider the case where a pilus can only generate a pulling force and no pushing force, thus the force is zero if *l*_c_ ≤ *l*_f_.

Experimentally, it has been shown that the pilus force affects the retraction velocity by

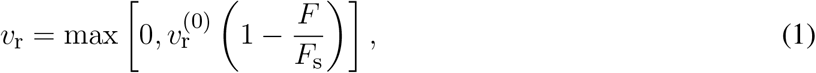

with the stalling force *F*_s_ [19]. The force of a pilus also affects the unbinding from the substrate. Pilus detachment is modelled by a Poisson process with the rate

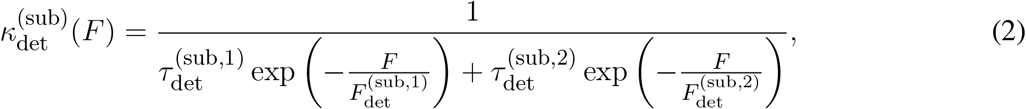

for detachment from the substrate, as motivated by [15]. Here, we introduce the detachment times 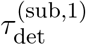 and 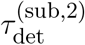 and the detachment forces 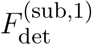 and 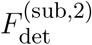. For detachment from a turbine disk, we describe the rate by a simpler relation

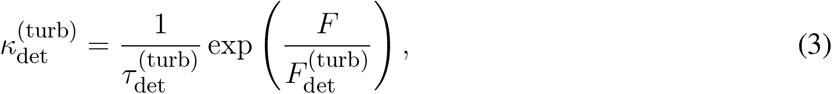

with the detachment time 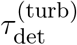 and detachment force 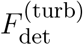.

### 2.3 Cell forces and motility

We model pili as Hookean springs with the spring constant *k*_p_. For a cell *i* at location 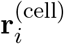, an attached pilus *j* causes a force 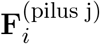 at the pilus anchor point 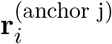 on the surface of the cell in the direction of the point 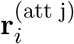 where the pilus tip is attached (see Fig. 1C).

Next to the active pili force, passive excluded volume forces are acting on the cells. Cells are located in a two-dimensional box with size *L* × *L* and a cell *i* that overlaps with the boundary wall is exposed to a repulsive force 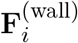 (see Fig. 1C). This force is modelled as a harmonic interaction with spring constant *k*^(wall)^ and acts on the cell center in the normal direction of the boundary wall if the overlap is smaller than the cell radius *R*. Additionally, intersections of two cells *i* and *k* lead to a repulsive force 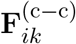 of the centers of both cells with the spring constant *k*^(c–c)^ (see Fig. 1C). A similar type of repulsive force 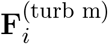 is acting between the cells and the turbine disks *m* with the spring constant *k*^(turb)^ (see Fig. 1C).

The total force of the cell *i* is given by

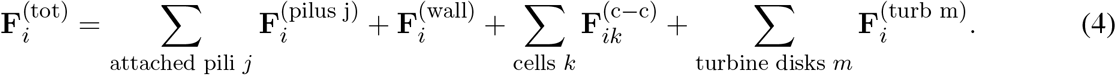

Additionally, the total torque acting on a cell is

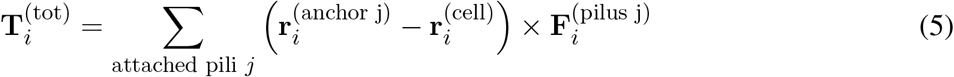

In the overdamped limit [26], a force mediates a translational motion of the cell with the velocity

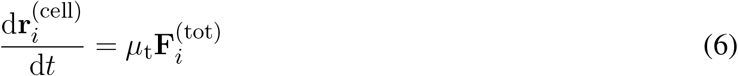

and the torque leads to a rotation with the angular velocity vector

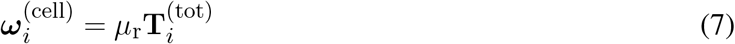

Here, we introduce the translational mobility *μ*_t_ and the rotational mobility *μ*_r_. The same forces and torques cause an equivalent displacement of the pili anchor points 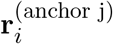 and attachment points 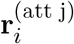. The mobilities introduced here are a result of friction with the viscous solvent and the substrate on which the cells are moving.

### 2.4 Turbine torque and rotation

The turbine is only able to undergo rotational motion. The total torque acting on the turbine is given by

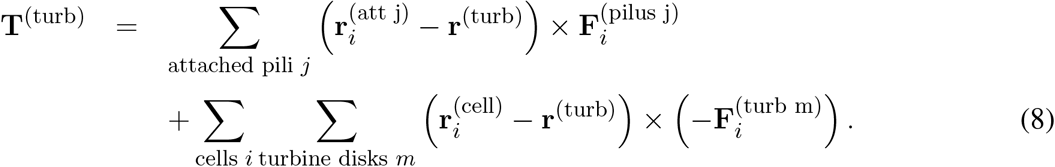

The rotation of the turbine is modelled in the overdamped limit with a mobility *μ*_turb_. In that case, the turbine angular velocity vector is

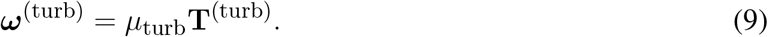

### 2.5 Parameters and details of numerical solution

The simulations were performed on the local computing cluster of the Max Planck Institute for the Physics of Complex Systems (Dresden, Germany), consisting of x86-64 GNU/Linux systems. The code was written in C++. We use an Euler algorithm to solve the equations of motion with a time step *δt*. While this is one of the most simple numerical schemes to solve the equations of our computational model, it often leads to numerical errors and instabilities when modelling molecular dynamics systems for long times [27]. We do not expect that this is a problem in our system due to the stochastic nature of the pili binding and unbinding, which basically represents our system as a series of many short time events, continuously interrupted by rearrangements in the pili network. Hence, we do not expect any differences in the qualitative outcome of the simulations.

If not stated otherwise, we use the of parameters provided in Table 1. Most parameters we use are based on previous studies. The excluded volume spring constants *k*^(c–c)^, *k*^(turb)^ and *k*^(wall)^ have no effect on the simulation outcome as long as they are chosen large enough to be able to compete with the pili forces. For the remaining parameters, e.g. the turbine mobility, we do not expect a qualitative difference in the results of the simulation.

**Table 1.**
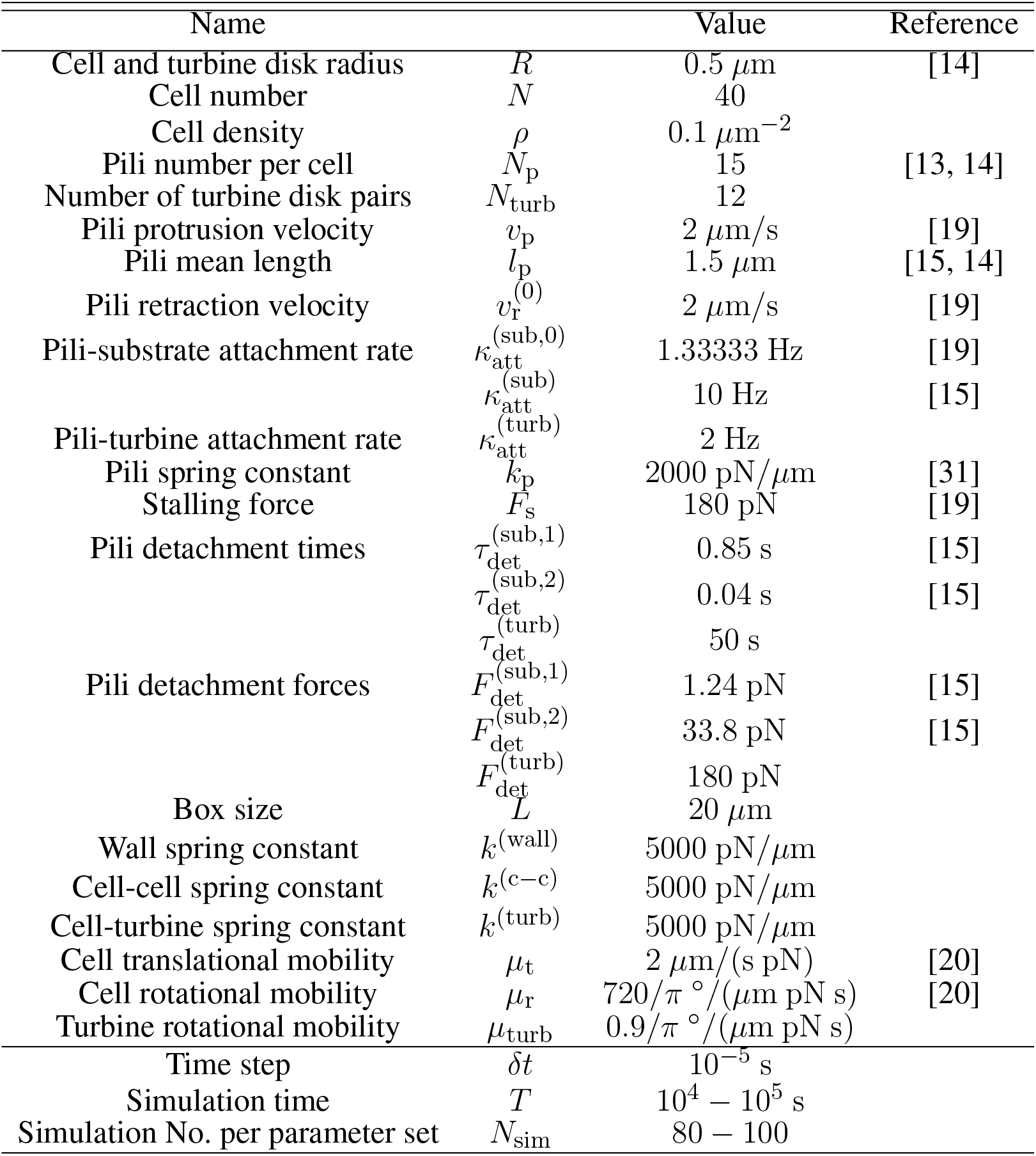
List of parameters used in this study.

We initialize the simulation by randomly distributing the cells in the box and only analyse the turbine rotation after an initialization period of 1000 s. To calculate the angular velocity of the turbine rotation, we compute

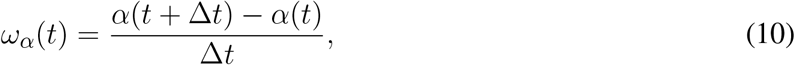

with ∆*t* = 0.5 s and the turbine orientation *α* (see Fig. 1B).

## 3 RESULTS

In the following, we first demonstrate that bacteria binding to the turbine with the help of pili can cause a persistent turbine rotation over time scales that exceeds the pili detachment times significantly. Next, we study how the persistent turbine rotation depends on the binding properties of the pili to the turbine and the number of cells in the system. This allows us to unravel the underlying mechanism that causes the persistent rotation of the turbine. Finally, we propose a system where an asymmetric binding of the pili to the turbine causes a permanent unidirectional rotation of the turbine.

### 3.1 Adhesion of motile bacteria drives turbine rotation

To investigate how the adhesion of cells to a turbine affects the turbine rotation, we first simulate the computational model for three different cases: (1) pili detach with detachment times of 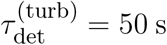 (see Fig. 2A and Movie S1), motivated by binding times in the order of a few minutes inferred from cell trajectories on plastic surfaces [14], (2) pili detach with 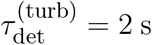 (see Movie S2), motivated by considerably smaller binding times experimentally measured for BSA coated beads [15] and (3) pili not bind to the turbine at all (see Movie S3) and the only way how cells interact with the turbine is by excluded volume forces. In Fig. 2B, we show the trajectories of the turbine angle *α*. We find that for the largest detachment time, 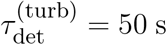, the rotation is the strongest. If pili are not permitted to attach to the turbine, the rotation appears to be the weakest. Additionally, the distribution of velocities shows that if pili cannot bind to the turbine, the turbine will move with smaller angular velocities *ω*, while the distributions seem to not depend on the detachment rate 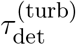, as long as pili are capable of binding to the turbine. To quantify how strong the rotation is, we compute the angular mean squared displacement (angular MSD), given by

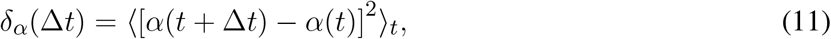

(see Fig. 2C). Indeed, the angular MSD is the highest for 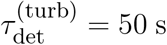 and the lowest for the case where pili do not attach to turbine disks. For 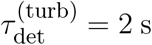, the angular MSD is identical to the case 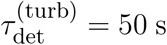 for small time differences (∆*t*). In both cases, 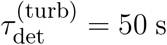 and non-attaching pili, the angular MSD follows a linear scaling, *δ_α_* ∝ ∆*t*, corresponding to a diffusive regime. For 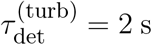, we find a superdiffusive regime for intermediate time differences ∆*t* ≈ 10 − 10^3^ s. For particles undergoing translational motion, such superdiffusive regimes usually emerge from ballistic motion. For the rotatable turbine, this correspond to a regime where the turbine moves persistently in one direction. Indeed, such a behaviour is observed in Fig. 2B where trajectories of the angle *α* move persistently in one direction over periods of hundreds to thousands of seconds before the rotation turns towards the opposite direction. For larger time differences, the angular MSD becomes again diffusive, implying that while the rotation is persistent over a certain time scale, for larger time intervals it becomes random again.

**Figure 2.**
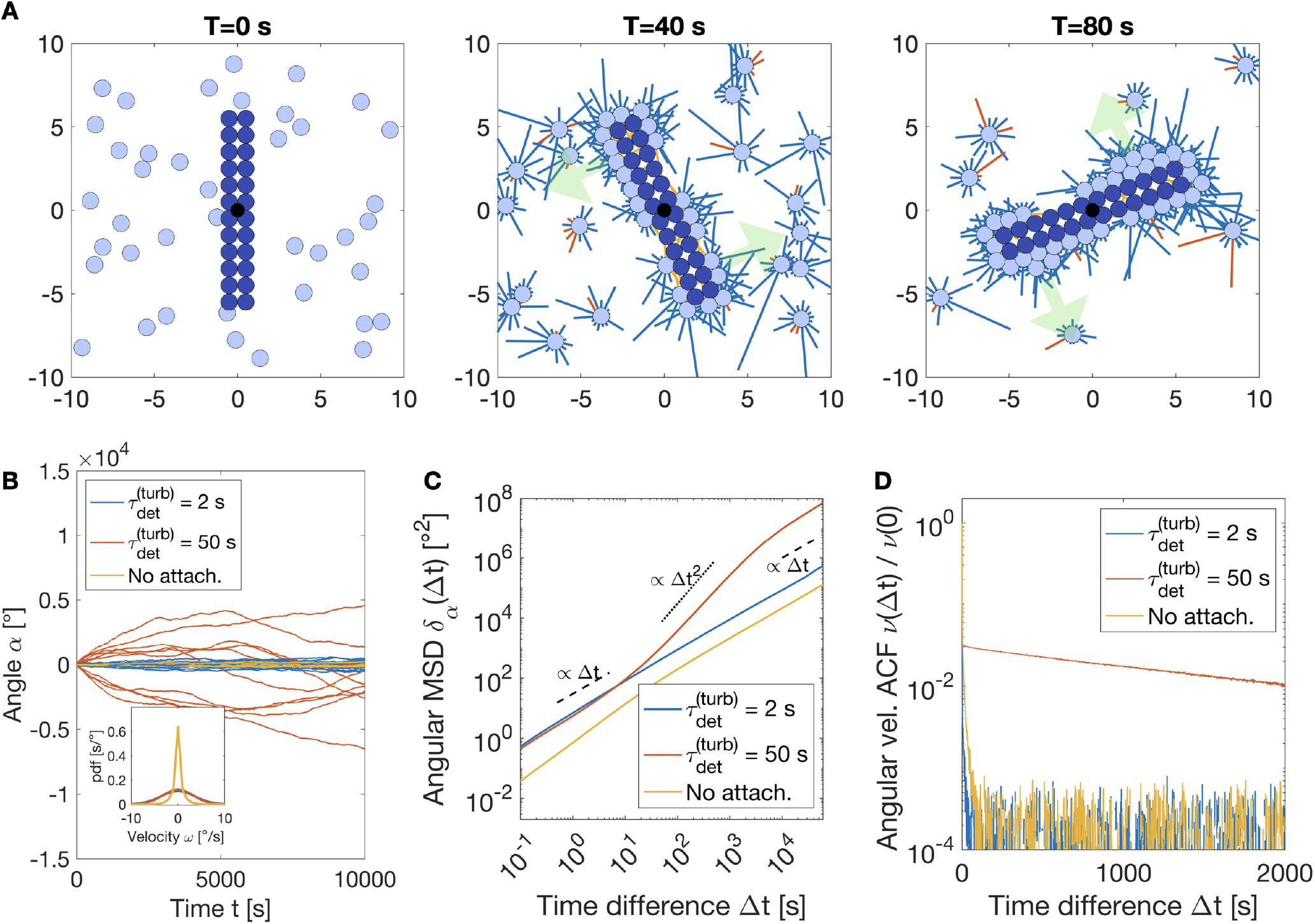
Analysis of rotary motion of a turbine in a bath of twitching bacteria. **(A)** Snapshots of the simulations of a turbine in a bath of bacteria with pili detachment rate 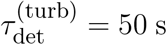. Blue pili are not attached, orange pili are attached to the turbine and red pili are attached to the substrate. The green arrow indicates the direction of the turbine rotation.**(B)** Turbine angle *α* as a function of the time *t* (see Fig. 1B.). The inset figure shows the probability density function of the turbine angular velocity *ω*. **(C)** Angular mean squared displacement (angular MSD) of the turbine angle, see Eq. (11). **(D)** Normalised angular velocity autocorrelation function of the turbine angle, see Eq. (12).

To learn more about how persistent the turbine rotation is, we compute the angular velocity autocorrelation function (angular velocity ACF), given by

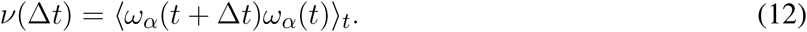

Independently of whether pili can bind to the turbine or not, we find that the correlation function *ν*(∆*t*) is decaying with time (see Fig. 2D). This confirms our previous observation for the angular MSD where we found that for very large time differences the turbine rotation is diffusive and no longer shows signs of persistence. We also find that for 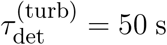, the normalised angular velocity ACF *ν*(∆*t*)/*ν*(0) has the slowest decay. This behaviour correlates with the persistent rotation for time differences between ∆*t* ≈ 10 − 10^3^. Surprisingly, we find that the decay of the angular velocity ACF is faster for 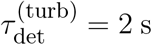 than for the case where pili do not bind to the turbine. We will provide an explanation for this behaviour in section 3.3. Before doing that, we will have a closer look at how the turbine rotation depends on the cell adhesion to the turbine disks and the number of cells in the system.

### 3.2 Turbine rotation is controlled by bacterial adhesion strength and cell number

We begin with a systematic analysis of how the angular MSD *δ_α_*(∆*t*) depends on the detachment time of pili from the turbine disks 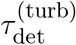, shown in Fig. 3A. For small time differences ∆*t* < 10 s, we find that *δ_α_* is independent of 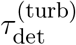 and exhibits a linear scaling, *δ_α_* ∝ ∆*t*, corresponding to a diffusive regime. For high enough values of the detachment time 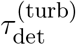 and ∆*t* > 10 s, the angular MSD becomes superdiffusive, *δ_α_* ∝ ∆*t*^2^. The duration of this superdiffusive regime increases with increasing 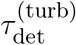 and is barely observable for detachment times 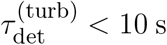. For even longer time differences ∆*t*, the MSD becomes diffusive again, *δ_α_* ∝ ∆*t*.

**Figure 3.**
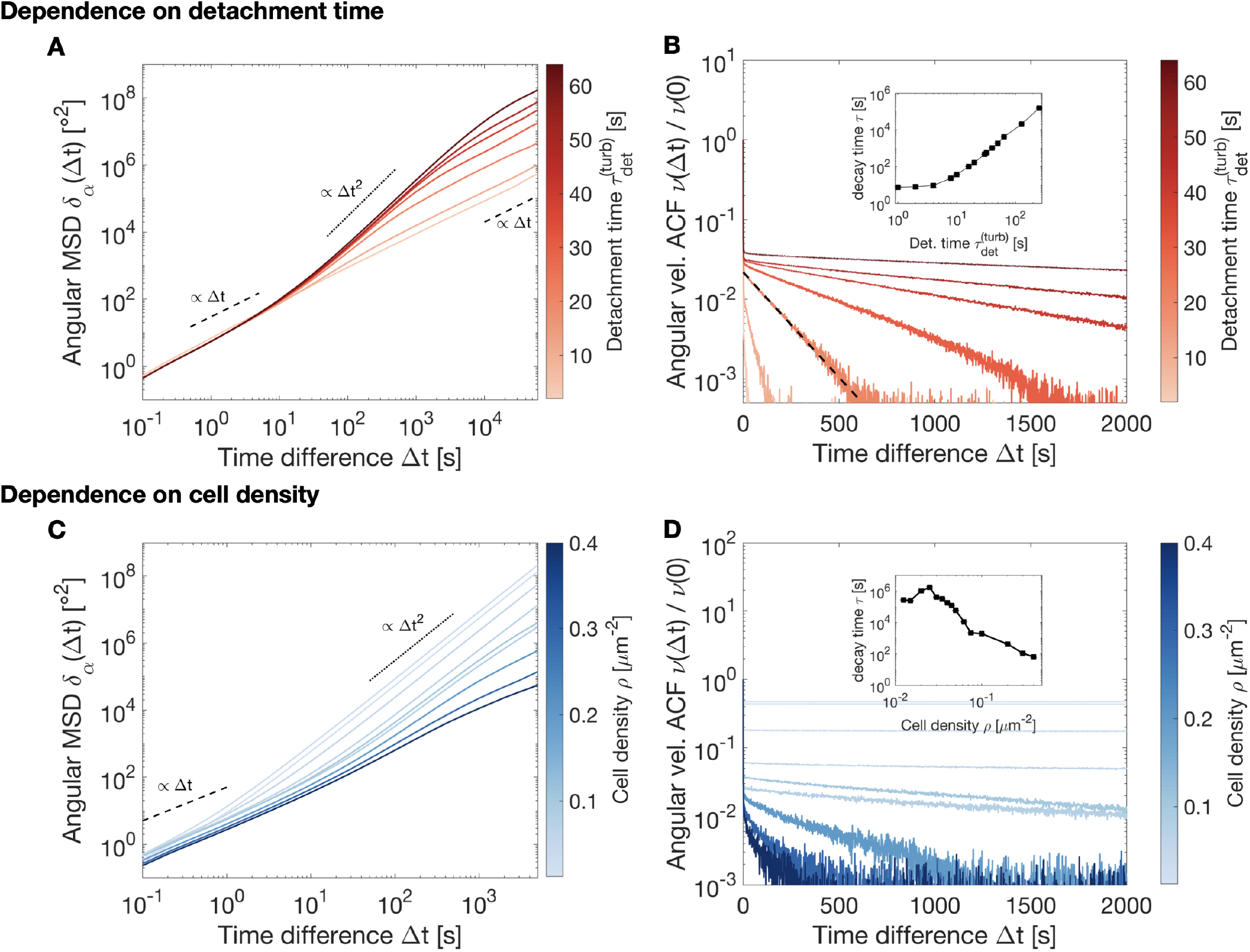
Dependence of the turbine rotation on pili detachment time 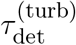 and cell density *ρ*. **(A and C)** Angular mean squared displacement (angular MSD) *δ_α_*(∆*t*) of the turbine angle *α* as a function of the pili-turbine detachment time 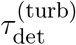 (with the cell density at *ρ* = 0.1 *μ*m^2^) or cell density *ρ* (with the detachment time 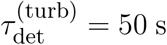). **(B and D)** Normalised angular velocity autocorrelation function (angular velocity ACF) *ν*(∆*t*)/*ν*(0) of the turbine angle *α* as a function of the pili-turbine detachment time 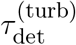 or cell density *ρ*. The inset figure shows the characteristic time scale of *ν*(∆*t*), resulting from a fit *ν*(∆*t*) exp (− ∆*t*/*τ*). To account for the initial drop of the angular velocity ACF, we only do the fitting for ∆*t* > 2s. An example fit is shown in **(B)** (black dashed line). Please note that for the investigation of the cell density dependence of the turbine rotation, we only consider a smaller range of ∆*t* to avoid too high numerical cost. As a result, we do not show the diffusive regime in the MSD for large values of ∆*t* since the angular velocity ACF is decaying with increasing ∆*t*.

Next, we investigate the normalised angular velocity ACF *ν*(∆*t*)/*ν*(0) (see Fig. 3B) and find that for ∆*t* < 2 s, it rapidly decreases. For ∆*t* ≥ 2 s, it exponentially decreases with the ∆*t*. By fitting a function of the form *ν*(∆*t*) ∝ exp (−∆*t*/*τ*) to this later regime, we identify the characteristic time *τ* of the exponential decay of *ν*(∆*t*) and can investigate its dependence on 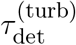. We find that it is increasing with increasing detachment times 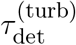. This is in accordance with the increasing duration of the superdiffusive regime of the angular MSD. The stronger pili of a cell bind to the turbine, the more persistently the turbine rotates in one direction before it inverts its direction.

Additionally, we investigated how the rotation of the turbine is affected by the density *ρ* of cells in the system. We find that with increasing density, the angular MSD *δ_α_* becomes smaller (see Fig. 3C). While we observe the diffusive regime of *δ_α_*(∆*t*) for ∆*t* < 10 s, a pronounced superdiffusive regime for larger ∆*t* vanishes if the cell density becomes too large. For the normalised angular velocity ACF *ν*(∆*t*)/*ν*(0) (see Fig. 3D), the characteristic decay time *τ* is initially increasing with cell density *ρ* and then decreases when *ρ* increases further.

In the following section, we provide qualitative arguments that explain the observed behaviours.

### 3.3 Unbinding dynamics of attached bacteria explains characteristic time of turbine rotation

To understand how the persistent rotation of the turbine in an otherwise symmetric system can emerge, we first consider how strongly cells bind to the turbine disks with the help of their pili and how pili binding affects substrate attachment of the remaining pili. Our hypothesis is the following: if cells stay attached to the same position on the turbine surface for an extended time, which is considerably larger than average attachment times of individual pili, the cells will continuously pull the turbine in the same direction. This mechanism is dramatically different to the previously reported rotary microdevices driven by swimming bacteria [4, 5], where cells collide with the microdevice and hence, push it. Instead, in our simulations, cells pull on the turbine.

In Fig. 4A, we show sketch of a cell which is attached to the turbine disk with some of its pili, while the other pili bind or unbind from the substrate. Only pili that protrude towards the turbine disks can attach, while pili protruding away from the turbine can only attach to the substrate. Thus, cells preferentially bind to the substrate in the direction away from the turbine. Since the cell is also attached with some of its pili to the turbine disks, it is pulling the turbine along the direction pointing from the turbine towards the cell. Thus, the cell pulls on the turbine, leading to a rotation of the turbine in the direction of the attached cell. This process is enhanced by the cooperation of multiple pili of a cell. To clarify this, we investigate the durations of cells attaching to the turbine disk before they detach again. To this aim, we solve a stochastic model of pili binding and unbinding, where a cell, possessing in total *N*_0_ pili that are all growing in the direction of the turbine, has *n* pili attached to the turbine. With a rate *k*_att_(*N*_0_ − *n*) a non-attached pilus attaches to the turbine, while with the rate *k*_det_*n* a pilus detaches from the turbine (see Fig. 4B). By describing the attachment and detachment as Poisson processes, we can numerically solve this system with the help of a Gillespie algorithm and investigate the mean time for a complete unbinding of all pili of a cell from the turbine (see Fig. 4C). We find that the mean unbinding time is increasing rapidly with the pili detachment time 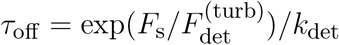, reaching around 10 s for *τ*_off_ = 2 s and 10^7^ s for *τ*_off_ = 50 s. For simplicity, we assume that all pili pull with their stalling force and that the detachment force is identical to the stalling force, 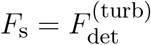. The duration of how long cells bind to the turbine corresponds to the duration of superdiffusive behaviour in Fig. 3A and also exhibits the same qualitative behaviour as the decay time of the angular velocity autocorrelation (see Fig. 3B). We do not expect a perfect quantitative agreement between the two time scales (Fig. 3B and Fig. 4C) as the simplified model of pili binding and unbinding of a cell to the turbine provided here ignores that cells can indeed move even if its pili are bound to the turbine, e.g. parallel to the turbine surface. Additionally, when multiple cells bind to the same region of the turbine, they will interact via excluded volume effects. This will lead to additional forces acting on the involved pili and thus, might enhance their detachment.

**Figure 4.**
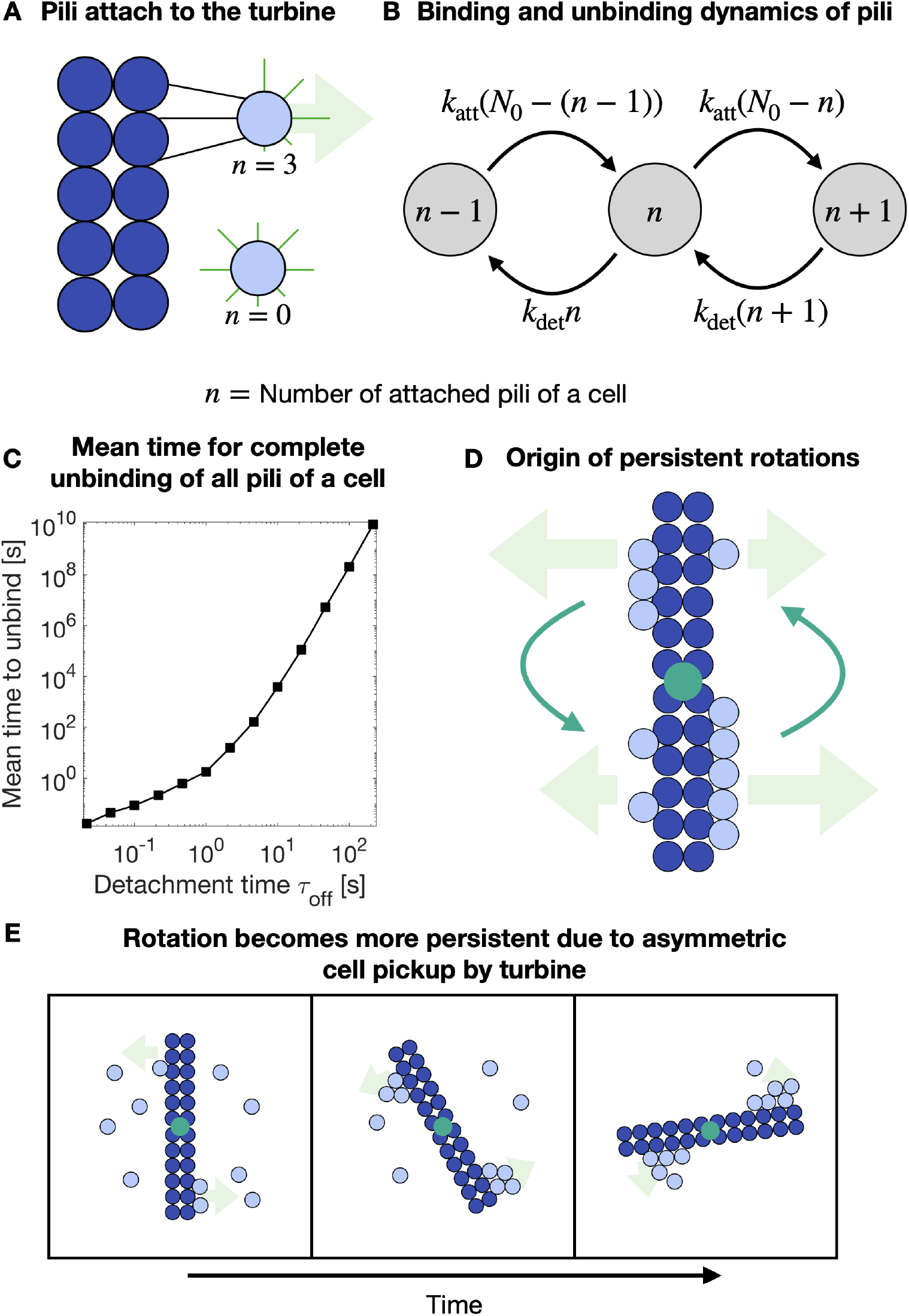
Overview of the mechanism driving the persistent rotation of the turbine. **(A)** A cell (light blue circle) with *n* = 3 or *n* = 0 pili attached to a turbine (dark blue circles). **(B)** Stochastic model of *n* pili attached to a turbine with pili binding (with constant rate *k*_att_) and unbinding (with constant rate *k*_det_). **(C)** Mean unbinding time to reach for the first time the state *n* = 0 in the stochastic model. Here, we chose *N*_0_ = 5, *n*(*t* = 0) = 5, *k*_att_ = 2 Hz, *k*_det_ = (exp 1)/*τ*_off_. **(D)** Sketch of how turbine rotation is affected by multiple cells binding to the turbine surface. The turbine rotates in the direction in which more cells are attached to its surface (turbine is pulled by cells). **(E)** Sketch of how turbine rotation leads to a pickup and binding of cells on the turbine surface.

We can now provide an explanation of the three time regimes observed in the angular MSD in Fig. 2C and angular velocity ACF in Fig 2D. For short times, ∆*t* < 10 s, individual pili stochastically bind and unbind to the substrate, pulling on the turbine, and additionally, cells randomly collide with the turbine. This causes small displacements of the turbine angle *α* in one direction, also explaining the very sudden drop of the angular velocity ACF for short times since the life time of such displacements is very small. This leads to random fluctuations of the turbine, and as a result, to a diffusive scaling. For larger times, 10 s < ∆*t* < 10^3^ s, asymmetries in the distribution of cells on the surface of the turbine lead to a unidirectional rotation of the turbine and as a result, to a superdiffusive regime. The persistence in the turbine rotation also explains the positive angular velocity ACF. The duration of this regime depends on how strong pili bind to the turbine disk (see Fig. 4C). Finally, for ∆*t* > 10^3^ s, the distribution of cells on the turbine surface is re-arranged up to such a degree that the turbine can change its direction of motion. This will then lead to a diffusive regime again. This time also corresponds to the characteristic time of the angular velocity ACF decay, confirming that for too large times, the turbine forgets its initial direction of motion.

Next, we consider the origin of the cell density dependent rotation of the turbine, see Fig. 3C and Fig. 3D. If the cell density is very small, often no cell will bind to the turbine most of the time and thus, the angular MSD is initially increasing with cell density. For moderate cell densities the difference between cells attaching to different sides of the turbine (see Fig. 4D) will be significant and due to the random asymmetry, the turbine will rotate persistently in one direction until cells randomly unbind from the turbine. If the cell density gets too large, more and more cells are pulling in the opposite direction of the turbine rotation, reducing its persistence.

There are additional processes that can have a significant impact on the turbine rotation: (1) Due to the rotation of the turbine, it will constantly “pick up” cells it collides with. For the traditional rotary microdevices that are driven by swimming bacteria that collide with the device and push it [4, 5, 6], this would lead to a torque that acts in the opposite direction of the original rotation. When cells pull on the turbine instead, the persistent rotation is being enhanced since asymmetric distribution of cells bound to the turbine disks is getting even stronger (see Fig. 4E). (2) In section 3.1, we found that the angular velocity ACF seems to decay faster for the case with pili attachment and 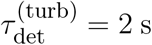, than for the case without pili attachment (see Fig. 2D). A possible explanation for this is that even without pili-turbine attachment, cells can still rotate the turbine due to collisions and resulting excluded volume forces. That way, we reproduce a system where cells push the turbine, instead of pulling on it. This process possesses its own characteristic time scales that are linked to the substrate motility of the cells.

To summarise, the persistent rotation of the turbine originates from persistent attachment of cells to the turbine over times scales much longer than the characteristic detachment times of individual pili. While the resulting rotation is indeed persistent and, depending on the binding properties, can go on over multiple revolutions, it is not permanently unidirectional. There is no asymmetry in the initial rotational direction of the turbine and even though the turbine can move in one direction over an extended time, this rotation will at some point reverse direction. Next, we will provide an example of how cells using type IV pili can drive a permanent unidirectional rotation of a turbine.

### 3.4 Permanent unidirectional rotation due to asymmetric cell-turbine attachment

In order to produce a permanent unidirectional rotation of the turbine, an asymmetry of the turbine is required. Typically, such assymetries are created by altering the turbine geometry [9, 5, 6], but here we have a chance to exploit asymmetries in the adhesion of cells to the turbine instead. In Fig. 5A, Fig. 5B and Movie S4, we provide an example of how altered pili binding properties on one side of the turbine wings can lead to a unidirectional rotation. Here, we consider different cases: (i) pili bind stronger to the manipulated turbine disks, corresponding to a larger detachment time 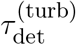 or (ii) pili bind weaker to the manipulated turbine disks, with a lower value of 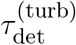. Depending on how pili bind to these regions, the direction of the turbine is affected as, on average, cells are more or less strongly adhered to the manipulated region and thus, more or less cells are bound to these sides of the turbine and mediate the turbine rotation. In case (i), the turbine rotates in the direction of manipulated region because more cells are attached there, for (ii) less cells are attached in the manipulated region and the turbine rotates in the opposite direction.

**Figure 5.**
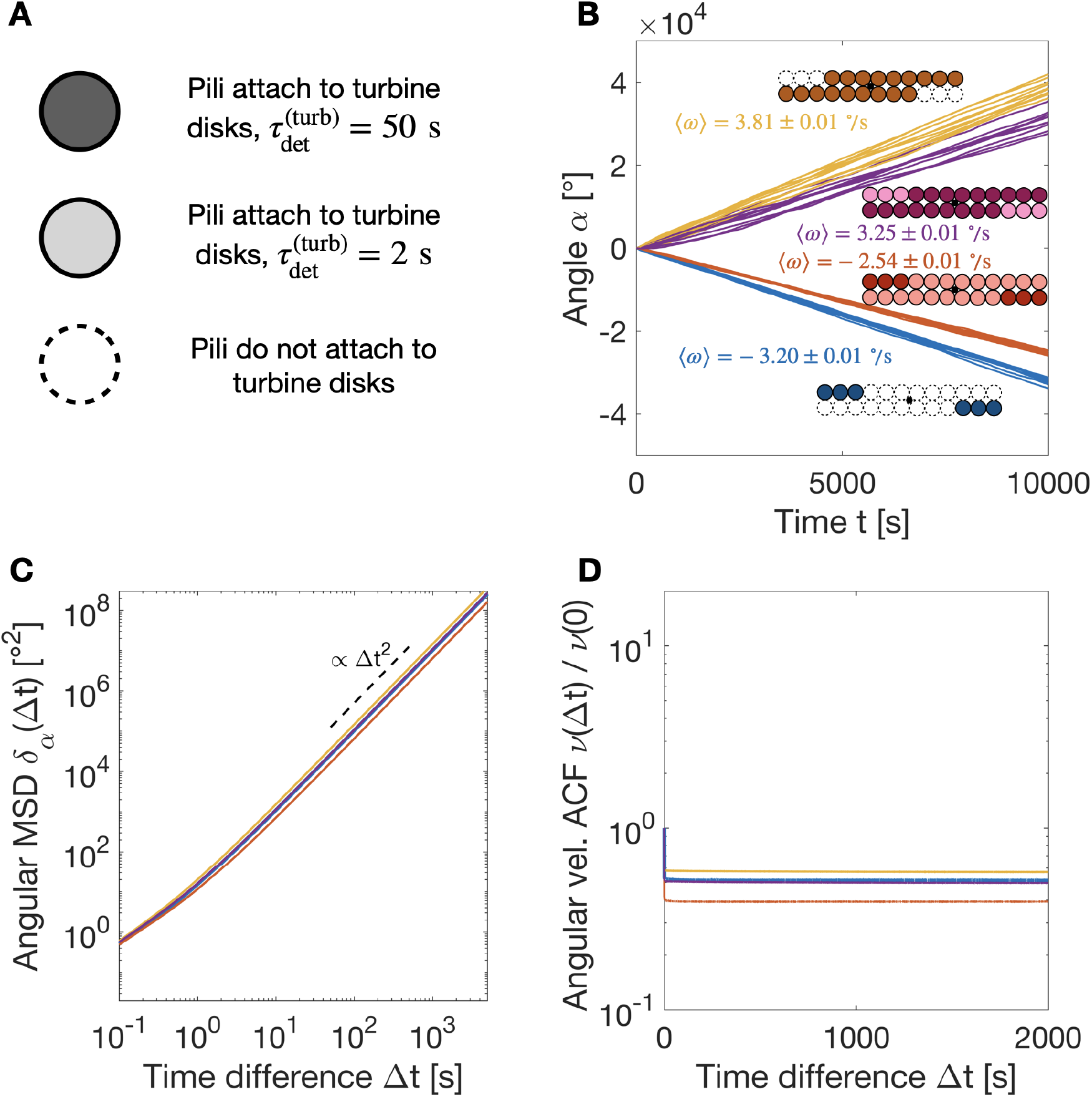
Unidirectional persistent turbine rotation is caused by asymmetric pili binding to the turbine. **(A)** Overview of the different types of turbine disks considered here. **(B)** Angle *α* of the turbine as a function of time *t* for different combinations of the manipulated turbine disks. We also provide the average angular velocity 〈*ω*〉 and its standard error for each condition. **(C)** The angular mean squared displacement exhibits superdiffusive (ballistic) scaling, *δ_α_* ∝ ∆*t*^2^. **(D)** The normalised angular velocity auto correlation function is no longer decaying with increasing time differences ∆*t*, corresponding to the unidirectional rotation.

We also estimate the average angular velocity *ω* of the turbine and find values around 2 − 4 ° /s. For the used rotational mobility of the turbine *μ*_turb_ (see Table 1), this corresponds to a turbine torque around 10 pN *μ*m, comparable to previously published values for rotary devices in bacterial baths of hundreds of swimming cells [5]. Here, however, the system consists of 40 cells only. Note that this is only the lower limit of the turbine torque and the same turbine might be able to generate larger torques if it experiences an opposing torque. In the absence of a counteracting torque, the rotation speed of the turbine is limited by the retraction velocity of pili, around 2 *μ*m/s [19].

We see evidence that the rotation is permanently unidirectional in the angular MSD (see Fig. 5C) which is superdiffusive (ballistic) for arbitrary time differences ∆*t*, equivalent to a rotation in one direction. Additionally, the normalised angular velocity ACF (see Fig. 5D) is no longer decaying with time ∆*t*, suggesting a constant rotation in the same direction.

In experiments, such manipulated regions on the surface of the turbine could be generated by coating it with a chemical such as BSA, which has been shown to alter the detachment time of the pili [28, 14, 15].

## 4 DISCUSSION

In this study, we investigated how bacteria that use type IV pili for surface motility may drive the rotation of a micron-sized turbine. We found that due to spatial asymmetries in the amount of cells that are bound to the turbine, a persistent rotation is observed over time scales that can strongly exceed the characteristic time scales of the pili. The persistent rotation is enhanced when pili bind more strongly to the turbine and weaker when the density of cells becomes larger. The observed persistent rotation of a symmetric turbine has a characteristic time scale. For larger times, the rotation direction is reversed stochastically and there is no preferred direction of rotation. A persistent and unidirectional rotation can be generated by altering the binding properties of the type IV pili on parts of the turbine in an asymmetric manner.

Here, we have shown that microdevices immersed in a bath of bacteria that use pili, such as *Neisseria gonorrhoeae* or *Pseudomonas aeruginosa*, can efficiently harness the power of the type IV pilus machinery. This system is particularly superior to previously reported microdevices driven by swimming bacteria [4, 5, 6], since the involved molecular motor pilT is the strongest known molecular motor [19] and a single cell is capable of generating forces in the nano-Newton range.

In the future, it will be interesting to investigate the interplay of asymmetries in the geometrical and adhesive properties of the turbine. Additionally, we expect that the binding of pili with the substrate will also have an effect on the turbine rotation. Higher values of the pili-substrate detachment times 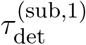 and 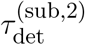 will lead to an increase in the torque exerted by attached cells on the turbine and will likely exceed the observed lower limit of the torque of around 10 pN *μ*m. This will enable the turbine to more efficiently harness the work generated by type IV pili. Importantly, one needs to consider that if pili bind too strongly to the substrate, their substrate mobility will be weakened [16] and hence, it will be less likely that cells will be in the vicinity of the turbine. Furthermore, cells pulling in the direction opposite to the turbine rotation will exert larger forces opposing the rotation. In the future, it will be interesting to investigate the optimal substrate binding properties to maximize the turbine torque. Additionally, bacteria use type IV pili to form aggregates consisting of hundreds to thousands of cells [12, 20, 21, 29, 30]. The impact of this aggregation process on the turbine rotation remains unclear.

## AUTHOR CONTRIBUTIONS

WP and VZ designed the research. WP carried out all numerical solutions. WP carried out analytical calculations. WP analyzed the data. WP and VZ wrote the article.

## DATA AVAILABILITY STATEMENT

The source code of the compuational model can be accessed via https://github.com/wolframponisch/Pili-Driven-Bacterial-Turbine.git. The Matlab scripts used to analyse the data and generate the figures are available upon request.

## REFERENCES

[1] Martel S, Tremblay CC, Ngakeng S, Langlois G. Controlled manipulation and actuation of micro-objects with magnetotactic bacteria. Applied Physics Letters 89 (2006) 233904.

[2] Behkam B, Sitti M. Effect of quantity and configuration of attached bacteria on bacterial propulsion of microbeads. Applied Physics Letters 93 (2008) 223901.

[3] Darnton N, Turner L, Breuer K, Berg HC. Moving fluid with bacterial carpets. Biophysical journal 86 (2004) 1863–1870.

[4] Hiratsuka Y, Miyata M, Tada T, Uyeda TQ. A microrotary motor powered by bacteria. Proceedings of the National Academy of Sciences 103 (2006) 13618–13623.

[5] Angelani L, Di Leonardo R, Ruocco G. Self-starting micromotors in a bacterial bath. Physical review letters 102 (2009) 048104.

[6] Di Leonardo R, Angelani L, Dell’Arciprete D, Ruocco G, Iebba V, Schippa S, et al. Bacterial ratchet motors. Proceedings of the National Academy of Sciences 107 (2010) 9541–9545.

[7] Pietzonka P, Fodor É, Lohrmann C, Cates ME, Seifert U. Autonomous engines driven by active matter: Energetics and design principles. Physical Review X 9 (2019) 041032.

[8] Wall D, Kaiser D. Type iv pili and cell motility. Molecular microbiology 32 (1999) 01–10.

[9] Harshey RM. Bacterial motility on a surface: many ways to a common goal. Annual Reviews in Microbiology 57 (2003) 249–273.

[10] Klausen M, Heydorn A, Ragas P, Lambertsen L, Aaes-Jørgensen A, Molin S, et al. Biofilm formation by pseudomonas aeruginosa wild type, flagella and type iv pili mutants. Molecular microbiology 48 (2003) 1511–1524.

[11] Imhaus AF, Duménil G. The number of neisseria meningitidis type iv pili determines host cell interaction. The EMBO journal 33 (2014) 1767–1783.

[12] Taktikos J, Lin YT, Stark H, Biais N, Zaburdaev V. Pili-induced clustering of n. gonorrhoeae bacteria. PLoS One 10 (2015) e0137661.

[13] Holz C, Opitz D, Greune L, Kurre R, Koomey M, Schmidt MA, et al. Multiple pilus motors cooperate for persistent bacterial movement in two dimensions. Physical review letters 104 (2010) 178104.

[14] Zaburdaev V, Biais N, Schmiedeberg M, Eriksson J, Jonsson AB, Sheetz MP, et al. Uncovering the mechanism of trapping and cell orientation during neisseria gonorrhoeae twitching motility. Biophysical journal 107 (2014) 1523–1531.

[15] Marathe R, Meel C, Schmidt NC, Dewenter L, Kurre R, Greune L, et al. Bacterial twitching motility is coordinated by a two-dimensional tug-of-war with directional memory. Nature communications 5 (2014) 1–10.

[16] Pönisch W, Weber CA, Zaburdaev V. How bacterial cells and colonies move on solid substrates. Physical Review E 99 (2019) 042419.

[17] Maier B, Potter L, So M, Seifert HS, Sheetz MP. Single pilus motor forces exceed 100 pn. Proceedings of the National Academy of Sciences 99 (2002) 16012–16017.

[18] Chattopadhyay S, Moldovan R, Yeung C, Wu X. Swimming efficiency of bacterium escherichiacoli. Proceedings of the National Academy of Sciences 103 (2006) 13712–13717.

[19] Maier B. The bacterial type iv pilus system–a tunable molecular motor. Soft Matter 9 (2013) 5667–5671.

[20] Pönisch W, Weber CA, Juckeland G, Biais N, Zaburdaev V. Multiscale modeling of bacterial colonies: how pili mediate the dynamics of single cells and cellular aggregates. New journal of physics 19 (2017) 015003.

[21] Pönisch W, Eckenrode KB, Alzurqa K, Nasrollahi H, Weber C, Zaburdaev V, et al. Pili mediated intercellular forces shape heterogeneous bacterial microcolonies prior to multicellular differentiation. Scientific reports 8 (2018) 1–10.

[22] Zhou K, Hennes M, Maier B, Gompper G, Sabass B. Non-equilibrium dynamics of growing bacterial colonies. arXiv preprint arXiv:2106.06729 (2021).

[23] Jin F, Conrad JC, Gibiansky ML, Wong GC. Bacteria use type-iv pili to slingshot on surfaces. Proceedings of the National Academy of Sciences 108 (2011) 12617–12622.

[24] Westling-Häggström B, Elmros T, Normark S, Winblad B. Growth pattern and cell division in neisseria gonorrhoeae. Journal of Bacteriology 129 (1977) 333–342.

[25] Chang YW, Rettberg LA, Treuner-Lange A, Iwasa J, Søgaard-Andersen L, Jensen GJ. Architecture of the type iva pilus machine. Science 351 (2016).

[26] Purcell EM. Life at low reynolds number. American journal of physics 45 (1977) 3–11.

[27] Bou-Rabee N. Time integrators for molecular dynamics. Entropy 16 (2014) 138–162.

[28] Whitchurch CB. Biogenesis and function of type iv pili in pseudomonas species. Pseudomonas (Springer) (2006), 139–188.

[29] Bonazzi D, Schiavo VL, Machata S, Djafer-Cherif I, Nivoit P, Manriquez V, et al. Intermittent pili-mediated forces fluidize neisseria meningitidis aggregates promoting vascular colonization. Cell 174 (2018) 143–155.

[30] Alston H, Parry AO, Voituriez R, Bertrand T. Intermittent attractive interactions lead to microphase separation in non-motile active matter. arXiv preprint arXiv:2201.04091 (2022).

[31] Biais N, Higashi DL, Brujić J, So M, Sheetz MP. Force-dependent polymorphism in type iv pili reveals hidden epitopes. Proceedings of the National Academy of Sciences 107 (2010) 11358–11363.

